# Absence of Local Retinotopy in the Mouse Optic Tract

**DOI:** 10.1101/2025.11.23.690078

**Authors:** Matteo Tripodi, Hiroki Asari

**Affiliations:** Epigenetics and Neurobiology Unit, EMBL Rome, European Molecular Biology Laboratory, Monterotondo, 00015, Italy; Scuola Internazionale Superiore di Studi Avanzati (SISSA), Trieste, 34136, Italy

**Keywords:** retinotopy, optic tract, retinal ganglion cells, lateral geniculate nucleus, mouse

## Abstract

Retinotopy is a fundamental organizational principle of the visual system, where neighboring neurons represent adjacent points in visual space. This spatial relationship is established by precise anatomical wiring across successive areas, e.g., from the retina to the lateral geniculate nucleus (LGN) to the visual cortex. To examine the precision of this topographic arrangement within the long-range projection axons themselves, we recorded retinal ganglion cell (RGC) axons in the mouse optic tract (OT) and mapped their receptive fields (RFs). As expected for a retinotopically organized area, we found that nearby LGN cell pairs had significantly smaller RF distances than distant pairs. In contrast, no such relationship was observed among RGC axons in the OT. Modelling analyses further confirmed that the observed RF distances in the OT were incompatible with any locally retinotopic arrangement. Instead, the OT retained only coarse topography, with ∼18° RF deviations or ∼40 µm axonal displacements from an ideal retinotopic organization. These results demonstrate that the mouse OT lacks fine-scale retinotopy and maintains only broad topographic structure.

## Introduction

The vertebrate visual system maintains spatial relationships from the retina to the downstream visual areas via topographic projections (Huberman et al., 2008; Seabrook et al., 2017; Cang et al., 2018). This retinotopic organization emerges early in development (Debski and Cline, 2002; McLaughlin and O’Leary, 2005; Cang and Feldheim, 2013), and provides a fundamental platform for visual processing, ensuring that nearby points in visual space are represented by neighboring neuronal populations. In mice, retinal ganglion cells (RGCs) project to >40 brain areas (Morin and Studholme 2014; Martersteck et al, 2017), with nearly all axons passing through the same pathway – the optic nerve, optic chiasm, and optic tract (OT) – before diverging to targets, such as the superior colliculus and the lateral geniculate nucleus (LGN). While retinotopy in these downstream areas has been well characterized (McLaughlin, et al., 2003; Piscopo et al., 2013; Molotkov et al., 2023), it remains elusive to what extent retinotopy is maintained in the long-range RGC axons themselves.

Previous anatomical and tracing studies have demonstrated that retinotopy is largely preserved in the optic nerve (Bunt and Horder, 1983; Reese, 2011; but see Horton et al. 1979). Axons arising from specific retinal quadrants occupy consistent positions within the optic nerve, forming a reliable mapping of the retinal geometry as they exit the eye. This topographic organization is, however, substantially disrupted in the optic chiasm (Marcus and Mason, 1995; Colerro and Guillery, 1998), where axons from the two eyes intersect and distribute themselves into either ipsilateral or contralateral side. The optic chiasm has thus been considered as a sorting hub, where retinotopy is relaxed to permit reorganization based on molecular cues, midline crossing decision, and eventual target-specific routing (Jeffery, 2001). After passing through the optic chiasm, RGC axons reorganize and partially recover retinotopic organization in the OT via pre-target sorting (Simon and O’Leary, 1991; Plas et al., 2005; Sitko et al., 2018). This indicates the presence of coarse retinotopy in the OT before reaching the target; however, questions remain on the precision and functional consequences of this reorganization.

Using in vivo electrophysiological recordings, here we mapped visual receptive fields (RFs) of individual RGC axons in the mouse OT. We quantified the spatial organization of these RFs, and took a data-driven modelling approach to evaluate how faithfully visual space is represented in the OT. Our results revealed a moderate degree of retinotopy, but not a fine-grained one. This highlights a robustness of the visual system to imprecision in long-range wiring, with refinement of the topographic organization taking place in each target area.

## Results

In a brain region with retinotopy, by definition, neighboring cells have adjacent receptive fields (RFs), representing nearby points in visual space (Cang and Feldheim, 2013; Seabrook et al., 2017). To examine if the optic tract (OT) has a retinotopic organization, we thus performed in vivo extracellular recordings of retinal ganglion cell (RGC) axons in the mouse OT, and mapped their RFs using white-noise stimuli and reverse correlation (e.g., **Fig. 1A-E**). As a positive control, recordings were also made from the dorsal lateral geniculate nucleus (LGN; e.g., **Fig. 1F-J**), a major retinorecipient area well-known to have a retinotopy (Piscopo, et al., 2013).

**Figure 1:**
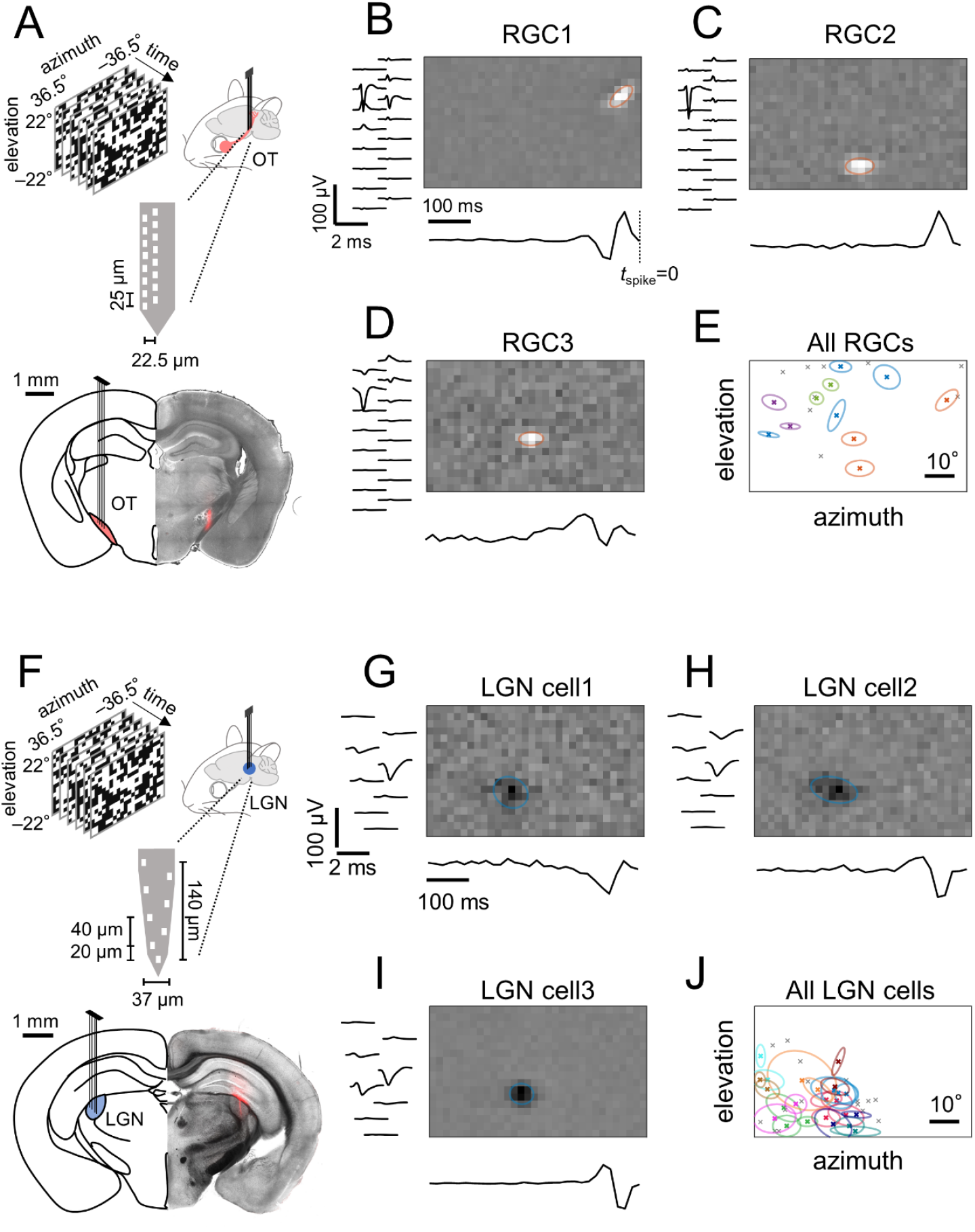
Receptive fields (RFs) of retinal ganglion cell (RGC) axons simultaneously recorded in the mouse optic tract (OT) are more widely spread across the visual field than those of lateral geniculate nucleus (LGN) cells. A: Schematic of OT recordings in awake head-fixed mice and a representative histological image showing the electrode position (DiI, red). B–D: Spatial RF (top) and temporal kernel (bottom) of three representative single-units that had the largest signals on the same recording site from a representative OT recording (left, average spike waveform in each recording site). Orange ellipses show a two-dimensional Gaussian envelope (at 1.5*σ*) fitted to the spatial RF profile of each unit. E: RF center locations of all simultaneously recorded single-units from the representative OT recording (crosses; N=20). Those with the largest signals on the same recording site are shown with the Gaussian envelopes of the same color. F–J: Corresponding data for a representative LGN recording (F, schematic and histological image; G–I, spatial RF, temporal kernel, and spike waveforms of three representative units that had the largest signals on the same recording site; J, RF center locations of all simultaneously recorded single-units, N=40).

Using multichannel silicon probes (4 shanks, each with 8-16 recording sites; **Fig. 1**), we simultaneously recorded 6±4 single-units from the OT (mean ± standard deviation, 33 animals) and 17±16 single-units from the LGN (17 animals). We found that the measured RFs of those single-units from the OT recordings were widely distributed across the visual stimulation area (73° and 44° in azimuth and elevation, respectively; RF size range, 1.3°–13.1°; e.g., **Fig. 1E**). This was also the case even for those units that had the largest signals on the same recording site, hence were supposedly located near each other in close proximity to the recording site (e.g., **Fig. 1B-D**). In contrast, the measured RFs of LGN cells were spatially more clustered (e.g., **Fig. 1J**) and demonstrated substantial overlap, especially among neighboring cells (e.g., **Fig. 1G-I**) as expected from known retinotopy (Huberman et al., 2008; Seabrook et al., 2017). Such qualitative comparison already suggested that the mouse OT lacks retinotopy.

For quantitative population-level data analysis, we examined how the RF distance relates to the physical distance between simultaneously recorded units (**Fig. 2**). We first used the electrode distance as a proxy of the distance between the units, and found a significant positive correlation for the LGN (Pearson correlation coefficient R=0.42, p<0.001, N=4491 pairs; **Fig. 2C**), but not for the OT (R=-0.01, p=0.8, N=756 pairs; **Fig. 2A**). The RF size differences between the pairs were small (LGN, 1.4°±0.9°; OT, 1.1°±0.6°; median ± median absolute deviation), indicating that consistent populations were sampled in each group. We next made a comparison between nearby and distant cell pairs, where nearby cell pairs were defined as those single-units that had the largest signals on the same recording site (i.e., zero electrode distance), whereas distant pairs as those with the largest signals on different recordings sites (i.e., non-zero electrode distance). In the LGN, the measured RF distance was significantly shorter for nearby cell pairs (4.8°±2.3°, median ± median absolute deviation; N=150 from 11 mice) than for distant cell pairs (14.0°±5.8°; N=4341 from 17 mice; p(U-test)<0.001; **Fig. 2D**). In the OT, however, RF distances did not differ between nearby (21.3°±15.1°, N=39 from 18 mice) and distant pairs (29.0°±12.7°, N=717 from 33 mice, **Fig. 2B**; p(U-test)=0.10). Thus, neighboring RGC axons in the mouse OT do not necessarily represent adjacent visual locations.

**Figure 2:**
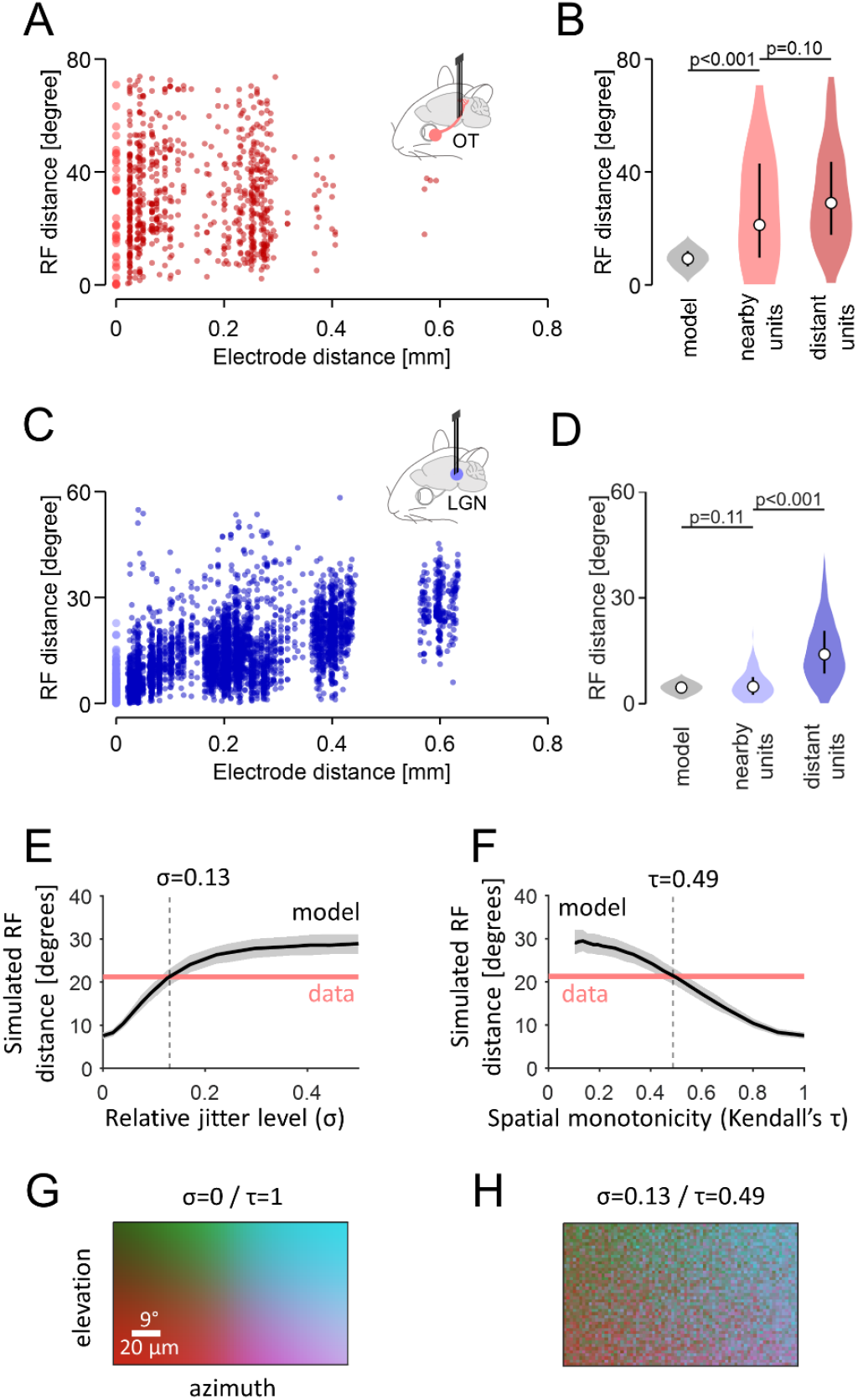
The mouse optic tract (OT) lacks local topographic organization, while maintaining global retinotopy. A,B: The RF center distance between RGC axons in the mouse OT as a function of the distance between the recording sites (A), and the probability distribution of the RF center distance (B: light red, nearby pairs with the largest signals on the same recording site, N=39 from 18 animals; dark red, distant pairs with the largest signals on different recording sites, N=717 from 33 animals; gray, expected distribution for nearby pairs by Monte Carlo simulation): circle, median; bar, the first and third quartiles. C,D: Corresponding data for LGN cells (light blue, N=150 nearby pairs from 11 animals; dark blue, N=4341 distant pairs from 17 animals; gray, Monte Carlo simulation). E,F: Simulated RF distance between nearby RGC axons at different retinotopy levels. Additive Gaussian noise (at different jitter levels, *σ*; E) was introduced to RF locations to achieve different spatial monotonicity levels (Kendall’s *τ*) of their topographic organization (F): black line, median; gray shade, 25 and 75 percentile range; black dotted line, expected *σ* and *τ* values given the measured data (red). G,H: Simulated topographic map (color-coded) of the mouse OT (G, precise retinotopy; H, moderate retinotopy corresponding to experimental data).

To further evaluate this, we modelled the expected RF distance for nearby cell pairs under the assumption of retinotopy (see Methods for details). For LGN, Monte Carlo simulations yielded an estimated RF distance between 4.2° and 4.9° (95% confidence interval; median, 4.6°; **Fig. 2D**), given the retinotopy gradient of 0.11°/µm (Piscopo et al., 2013) and a reliable single-unit recording range of 40 µm from an extracellular electrode (Anastassiou et al., 2015). This is consistent with the measured RF distance (median, 4.8°), validating our modelling approach. Applying the same logic to the mouse OT (diameter, ∼300 µm; visual field extending ∼135° in both azimuth and elevation; Paxinos and Franklin, 2001), our model predicted the median RF distance of 7.9°–10.7° for nearby pairs (median, 9.3°; **Fig. 2B**) if RGC axons were arranged retinotopically. This disagreed with the measured RF distance (median, 21.3°), hence rejecting the presence of retinotopy imposed as a key premise of the model.

To what extent does the mouse OT retain topographic organization? To address this question, we extended our model by introducing additive Gaussian noise to the RF location of the simulated RGC axons (see Methods for details), where *σ*=1 corresponds to the represented visual field (135°). We then used Kendall’s *τ* as a measure of spatial monotonicity to quantify the degree of retinotopy, where *τ*=1 and 0 represent precise and no retinotopy, respectively. As expected, the retinotopy level *τ* decreased with increasing noise level *σ*, and the estimated RF distance of nearby RGC axons expanded (**Fig. 2E,F**). From the intersection between the measured and simulated RF distances, we then identified that the mouse OT had a moderate level of retinotopy (*τ*=0.49) with *σ*=0.13, corresponding to a deviation of RF location from a retinotopically ideal position by ∼18°, or equivalently, a displacement of RGC axons by ∼40 µm (**Fig. 2G,H**). This indicates that the mouse OT lacks local retinotopy, while maintaining topographic organization at a global level.

## Discussion

Here we provide in vivo electrophysiological evidence for the absence of local retinotopy in the mouse optic tract (OT). Both population-level analyses and data-driven models failed to identify a fine-scale topographic organization in the OT that would make neighboring retinal ganglion cell (RGC) axons encode adjacent locations in the visual field. In contrast, corresponding analyses on the dorsal lateral geniculate nucleus (LGN) supported the presence of retinotopy as expected from previous studies (Huberman et al., 2008; Seabrook et al., 2017), thereby serving as a control for our functional circuit characterization.

We acknowledge several caveats in our results. First, the OT is predominantly comprised of RGC axons, but also contains axons from other sources, such as the parabigeminal nucleus (Reinhard et al., 2019; Tokuoka et al., 2020) and the superior colliculus (Gale and Murphy, 2014). Our physiological criteria (i.e., robust visual responses with short latencies and well-defined, spatially-confined linear RFs) may not be sufficient to fully exclude those nontargets. Second, while the OT is largely intermingled to form a broad retinotopy, there is partial and graded segregation of RGC axons by the cell type and projection target (Erskine and Herrera, 2014; Robles et al., 2014). Our models did not incorporate such typology, and thus may underestimate the actual degree of retinotopy in OT. Another caveat is that the nodes of Ranvier from adjacent RGC axons can be offset by up to ∼70 µm (Butt et al., 1994); hence, they may not be detected on the same recording site. Nevertheless, the units we defined as “nearby” should still be in close proximity to a given recording site, and thus near each other if not the closest. This guarantees the validity and robustness of our data/model analyses.

Our finding that RGC axons in the mouse OT retain only coarse retinotopy implies a fundamental constraint of early visual pathway organization: i.e., pre-target axon sorting cannot establish precise spatial relationships over long distances (Plas et al., 2005). This is consistent with known limits of axon guidance mechanisms, such as Eph/ephrin signaling and axon fasciculation, that establish broad topographic gradients but lack the resolution required for single-axon precision (McLaughlin and O’Leary, 2005; Cang and Feldheim, 2013). Thus, fine retinotopy must be re-established within each target via activity-dependent mechanisms—including retinal waves and Hebbian plasticity—to impose local precision on coarse inputs (Debski and Cline, 2002; McLaughlin et al., 2003).

Such a two-step architecture likely reflects an adaptive circuit organization principle: by relaxing long-range wiring constraints, the system reduces developmental cost while increasing robustness. Small positional deviations in the OT have then minimal impact because final mappings are refined locally in individual targets. Moreover, each area remains free to optimize the retinotopic map for its own computational needs (Knapen, 2021), rather than relying on the geometric arrangement of incoming axons themselves. Testing these functional implications in the visual system—and examining if they extend across the central nervous system—will be an important direction for future studies.

## Materials and Methods

No statistical method was used to predetermine the sample size. The significance level was 0.05 in all analyses unless otherwise noted. All experiments were performed under the license 233/2017-PR and 220/2024-PR from the Italian Ministry of Health, following protocols approved by the Institutional Animal Care and Use Committee at European Molecular Biology Laboratory. The data analyses were done in Python and Matlab.

### Animals

Animals were housed on a 12h light-dark cycle, with ad libitum access to water and food. In total, 50 female wild-type mice (C57BL6/J; RRID:IMSR_JAX:000664), 4-19 weeks old (median, 9.2 weeks old) at the time of surgery, were used for in vivo electrophysiology (optic tract, 33 animals; lateral geniculate nucleus, 17 animals).

### In vivo electrophysiology

In vivo electrophysiology was performed as described previously (Tripodi and Asari, 2025). Briefly, we first implanted a head-plate to animals for fixing their head during in vivo electrophysiological recordings. Before the surgery, animals were injected with Carprofen (5 mg/kg) and then anaesthetized with isoflurane (4% for induction, 1% for maintenance in O_2_). During the surgery, the animals were placed in a stereotaxic frame (Stoelting 51625) with a heating pad (Supertech Physiological) to keep their temperature stable at 37°C; and ointment (VitA-Pos, Ursapharm) was applied on both eyes to prevent them from drying. A portion of the scalp was removed to expose the skull, and the periosteum was scraped away with a round scalpel to increase adherence of the dental cement. A titanium head-plate with a hole (diameter, 8mm) was then cemented on the skull with a mixture of cyanoacrylate (Loctite 401, Henkel) and dental cement (Paladur, Kulzer).

The skull surface was then glazed with a thick layer of cyanoacrylate to support the skull with mechanical, atmospheric and biological protection, while still allowing for visual identification of reference points (bregma and lambda). After the surgery, the animals were placed on a heating pad for recovery, and then housed in individual cages. During the following seven days, the mice were administered with analgesia (Carprofen; 50 mg/mL) diluted in drinking water. The animals were then rehoused together to reduce post-surgical isolation.

After recovery from the surgery, the animals were habituated to head fixation on the experimental apparatus for one week (twice a day for up to two hours). On the day of the recording, we first placed the subject animal in the recording set up with its head fixed, and determined the electrode penetration path to the target area (optic tract, [-1.34, +1.87, +4.74], [-1.70, +1.87, +4.74], or [-1.82, +2.35, +4.07] in [anterior-posterior (AP), medial-lateral (ML), dorsal-ventral (DV)] coordinates; LGN, [-2.3, +2.3, +2.8]) using the robotic stereotaxic system (StereoDrive, NeuroStar). The animal was then briefly anesthetized with isoflurane for about 5 minutes, and a hole was drilled around the electrode entry point on the skull. After the removal of the anesthesia, an acute silicon probe (P2, Cambridge Neurotech, **Fig. 1A**; or Buzsaki32L, Neuronexus, **Fig. 1F**) coated with a fluorescent dye (DiI stain, Invitrogen, D282) was lowered at 5 µm/s using the robotic arm until visual responses were found in the target area. A battery of visual stimuli (see below for details) were then presented for recordings.

After the recording session, the electrode position was verified histologically. After retracting the silicon probe, the mice were anesthetized (2.5% Avertin, 16 μL/g, intraperitoneal injection) and perfused with paraformaldehyde (PFA; 4% in phosphate buffer solution), followed by brain tissue harvest and overnight post-fixation in 4% PFA at 4°C. Coronal sections of the brain tissue (thickness, 100-150 μm) were then examined under a fluorescence microscope (Leica, LMD7000 with N2.1 filter cube) to visualize the trace left by the DiI stain on the probe (**Fig. 1A,F**).

### Visual stimulation

Visual stimuli were presented as described before (Boissonnet et al., 2023). Briefly, visual stimuli were projected to a spherical screen (radius, 20 cm) placed ∼20 cm from the animal’s left eye, covering about a quarter of its visual field (±36.5° and ±22° in azimuth and elevation, respectively).

A gamma-corrected digital light processing device (Texas Instruments, DLPDLCR3010EVM-LC) was used as a light source, where the green and the red light-emitting diodes (LEDs) were replaced with ultraviolet (UV; 365nm, LZ1-00UV00, LED Engine) and infrared (IR; 950nm, SFH 4725S, Osram) LEDs, respectively. The UV and remaining blue channels were used for visual stimulation (frame rate, 60 Hz; maximum intensity, 31 mW/m^2^), while the IR signals were recorded with a photodiode (PDA100A2, Thorlabs) for data synchronization. For receptive field (RF) mapping, a black-and-white binary noise stimulus was presented for 15 minutes, consisting of a 32-by-18 pixels checkerboard patterns, where each pixel randomly and independently flickered at 60 Hz while keeping the overall luminance of each frame constant at the mean intensity.

### Data analysis

We adapted previously established methods of spike sorting and data analysis (Boissonnet et al., 2023; Tripodi and Asari, 2025). In brief, we used Kilosort 2.0 to sort spikes with a set of default parameters, except for the spike detection threshold to be 6 during optimization. Single-units were then identified by clustering in principal component space using Phy for visualization and manual data curation. Only those units that maintained the average spike waveforms and autocorrelograms with a minimal refractory period of 1 ms were kept for subsequent analyses.

For the visual RF analysis, we employed stimulus ensemble statistical techniques (reverse correlation methods; 500 ms window; Δt = 1/60 s bin size). Specifically, we first obtained the linear spatiotemporal RF of each recorded cell by calculating a spike-triggered average (STA) of the “checkerboard” stimuli with ±1 being “white” and “black” for each pixel, respectively. As a quality measure, *p*-value was computed for each voxel against a null hypothesis that the STA follows a normal distribution *N*(0, 1/*C*), where *C* is the total number of spikes. For those with *p* < 10^-5^ at any voxel, we ran a singular value decomposition to obtain temporal and spatial filters, respectively, assuming the separability of RGC/LGN spatiotemporal RFs. We then fitted a two-dimensional Gaussian envelope to those spatial filters with a single, distinct and localized feature; and considered the center of the Gaussian as the RF center location (**Fig. 1**). Single-units with little or no visual responses were excluded as nontarget cells, such as the axons from the parabigeminal nucleus in the OT (Reinhard et al., 2019; Tokuoka et al., 2020). For the OT recordings, we also applied an RF size threshold of 15° to exclude axons from wide-field cells in the superior colliculus (Gale and Murphy, 2014; Relota et al., 2025). In total, we obtained 60 ON RGCs (peak latency of the temporal filter, 46±5 ms, median ± median absolute deviation; RF size, 3.6°±0.9°, estimated as the mean of the fitted Gaussian envelope short- and long-axis diameters at 1 *σ*), 133 OFF RGCs (51±8 ms; 3.5°±0.9°), 137 ON LGN cells (46±4 ms; 4.7°±1.2°), and 155 OFF LGN cells (46±5 ms; 4.3°±1.0°).

To a first approximation, at short distances, the amplitude of extracellular signals is inversely proportional to the distance between the cell and the probe (Anastassiou et al., 2015). We thus considered that the recorded units were physically located nearby if they both had the largest spike waveforms on the same recording site on the probe, whereas distant if they had the largest spike waveforms on different recording sites. Using these criteria, we identified pairs of nearby and distant single-units that were recorded simultaneously (optic tract, 39 and 717 pairs in total, respectively; LGN, 150 and 4341 pairs in total, respectively), and compared the distance between their RF center locations at the population level using Mann-Whitney U-test (**Fig. 2**).

### Model analysis

We used Monte Carlo methods to obtain an expected RF distance distribution of nearby cell pairs, given a retinotopy gradient *g*. In particular, we first randomly sampled two points in a sphere, where the radius *r* represents the maximum distance from an extracellular electrode for reliable single-unit recording. If the distance between the two points, *D*, was larger than an exclusion threshold *d* due to a physical mass of neurons, we then estimated the RF distance between two neurons located at those two sampled points as *D×g*. We repeated the procedure 1000*nn* times to compute the probability distribution function of the estimated RF distance, as well as a 95% confidence interval of the median value (of sample size *n*). For the mouse OT (*n* = 42; **Fig. 2A**), we used *r* = 20 µm for recording range, *d* = 2 µm for RGC axon diameter, and *g* = (represented visual field) / (optic tract diameter) = 135° / 300 µm (Paxinos and Franklin, 2001). For the mouse LGN (*n* = 150; **Fig. 2C**), we used *r* = 40 µm for recording range, *d* = 10 µm for cell body size, and *g* = 0.11°/µm (Piscopo et al., 2013).

To run simulations at different retinotopy levels for the mouse OT (**Fig. 2E-H**), we added noise to the simulated RF locations, drawn from a Gaussian distribution *N*(0, *σ*^2^) where *σ*=1 corresponds to the full range of the represented visual field (135°), or equivalently, the full width of the mouse OT (300 µm; Paxinos and Franklin, 2001). Simulations were done with *σ*=[0, 0.5] in steps of 0.05, as described above except that we randomly sampled two points within the distance *d* from a randomly chosen probe position within the visual stimulation area (73° and 44° in azimuth and elevation, respectively), and eliminated a trial if any cell on those two sampled points had its RF outside the visual stimulation area due to the added jitter. With increasing jitter, the simulated RF distance increased and the topographic map exhibited greater disorganization. To quantify the level of retinotopy, we used Kendall’s *τ* as a measure of spatial monotonicity, where *τ*=0 indicates a random mapping with no retinotopy; and *τ*=1 represents a perfect retinotopy. The intersection between the measured and the simulated RF distances was then used to estimate the level of retinotopic organization in the mouse OT.

## Acronyms

AP: anterior-posterior
DV: dorsal-ventral
IR: infrared
LED: light emitting diode
LGN: lateral geniculate nucleus
ML: medial-lateral
OT: optic tract
PFA: paraformaldehyde
RF: receptive field
RGC: retinal ganglion cell
STA: spike-triggered average
UV: ultraviolet.

## Acknowledgements

This work was supported by research grants from EMBL (H.A.). The EMBL Light Imaging Facility is acknowledged for support in histological image acquisition; EMBL IT Support for provision of computer and data storage servers; and the LAR facility for taking care of animals. We thank all the Asari lab members for useful discussions.

